# Preliminary analysis of a neotropical plant extract antiviral potential against *Chikungunya* and *Mayaro* viruses

**DOI:** 10.1101/2021.08.04.455105

**Authors:** Ellen Caroline Feitoza Pires, Francini Pereira da Silva, Karoline Schallenberger, Bruna Saraiva Hermann, Larissa Mallmann, Alini Godinho dos Santos, Wellington S. Moura, Eugênio E. de Oliveira, Guy Smagghe, Sergio Donizeti Ascêncio, Robson dos Santos Barbosa, Ilsamar Mendes, Juliane Deise Fleck, Bergmann Morais Ribeiro, Raimundo Wagner de Souza Aguiar

## Abstract

Chikungunya and Mayaro fevers are viral infectious diseases, without vaccine or treatment, that causes fever and arthralgia. Establishing novel antiviral tools capable of preventing or treating *Chikungunya virus* (CHIKV), and *Mayaro virus* (MAYV) infections are needed. The use of plant-based compounds that affect the replication cycles of these viruses has been proposed as a promising strategy. *Chiococca alba* (L.) Hitchc. is a neotropical plant used by Yucatec Mayas traditional healers, mainly, as antipyretic and antirheumatic. To evaluate the potential of *C. alba* methanolic extracts against CHIKV and MAYV through preliminary analysis *in vitro* and *in silico*. The cytotoxicity profile of two *C. alba* roots methanolic extracts in Vero cells was performed by lysosomal viability using the neutral red assay, and the antiviral potential was determined by plaque assay. We further assessed, through *in silico* computational predictions, the possible interactions between the active site of the nsP2 proteases of these viruses with some secondary metabolites present in *C. alba* extracts, identified by High-Performance Liquid Chromatography (HPLC). Our partial phytochemical analysis revealed the presence of flavonoids, and phenolic acids in the *C. alba* extracts. Our *in vitro* assays showed that both *C. alba* extracts inhibited more than 70% of CHIKV and MAYV activities at 60 µg/mL concentration. Based on our *in silico* computational predictions, the flavonoids naringin and vitexin showed the greatest affinity energies with the CHIKV and MAYV nsP2 proteases, revealing the great potential of these compounds as viral inhibitors. The findings described here indicates that *C. alba* extracts, or their secondary metabolites, as a potential source of novel antiviral compounds.

## Introduction

Arboviruses diseases have a high impact on public health in many countries of the neotropical regions, due to frequent disease outbreaks and epidemics [1]. *Chikungunya virus* (CHIKV) and *Mayaro virus* (MAYV) are arboviruses transmitted through female mosquito bite of the genus *Aedes* [2]. They belong to the *Togaviridae* family and the *Alphavirus* genus, and clinically, the disease caused by CHIKV and MAYV is similar to dengue, characterized by fever, headache, and arthralgia [3,4].

*Alphavirus* infects cells through membrane receptor-mediated endocytosis [5]. Their genomes have two Opening Reading Frames (ORF’s), the first encodes four non-structural proteins (nsP1, nsP2, nsP3, and nsP4), and the second encodes proteins related to the structure of the viral particle: capsid (C), envelope glycoproteins (E1 and E2) and two cleavage products (E3 and 6k) [6,7]. *Alphavirus* non-structural proteins are involved in viral replication and transcription and, therefore, are potential targets for the development of inhibitors, mainly nsP2 due to their multi-action in viral replication [8, 9].

Plant-derived substances have been widely studied as sources of compounds with antiviral activity [10]. Among these, we can mention the products of secondary metabolism of different plant species [11], which have shown antiviral potential against several viruses [12]. Examples of these metabolites are alkaloids, saponins, flavonoids, and coumarins [13]. However, besides having high antiviral activity, metabolites also need to have low cytotoxicity to cells. Among the metabolites with antiviral activity mentioned in the literature, flavonoids stand out [14].

Rubiaceae family plant species are known to produce secondary metabolites with pharmacological activities, mainly, flavonoids, alkaloids and saponins [15, 16, 17]. The species *Chiococca alba* (L.) Hitchc. (Rubiaceae) is a 2 to 3 meters high shrub endemic to the American continent, commonly known in Brazil as “Cainca” [18, 19]. The *C. alba* was used for medicinal purposed by the Yucatec Mayas traditional healers of Southern Belize, in Central America, to treat fever, cold, and muscle pains [20, 21]. Roots’ infusion is used in traditional medicine as antirheumatic, antiasthmatic, diuretic, anti-inflammatory, and antimicrobial activity [22, 23].

Considering the symptoms of intermittent fever caused by infection with CHIKV and MAYV, associated with the Mayas Yucatec ethnobotanical knowledge, this study aimed to evaluate the effectiveness of methanolic plant extracts from the *C. alba* roots, as a potential antiviral agent against CHIKV and MAYV in Vero cell lineage *in vitro*. We also analyzed the *in silico* interaction of two identified compounds in the plant extract with the CHIKV and MAYV nsP2 protease.

## Materials and methods

### Plant material

*C. alba* roots were collected in Formoso do Araguaia (11 ° 47’48 “latitude S, 49 ° 31’44” longitude O), Tocantins state, Brazil [24]. The plant was identified at the herbarium of the Department of Environmental Studies of the Federal University of Tocantins (Campus Porto Nacional), where the specimen voucher was deposited under the code (HTO-11.160). The research was authorized by the Sistema Nacional de Gestão do Patrimônio Genético e do Conhecimento Tradicional Associado – SISGEN, n° A77A809.

### Preparation of extracts

The plant material powder was subjected to two different extraction processes: by maceration (CAH21) and to the Soxhlet extractor (Marconi, model MA-487/6/25, Brazil) (CAH24). For the CAH21 process, powder roots (500g) were extracted at room temperature (25 °C) for three days with methanol (MeOH), and for the CAH24 process, powder roots (500g) were extracted at 60 °C for 4 h with MeOH. The samples were filtered through Whatman No. 1 filter paper (GE Healthcare Life Sciences, USA), and the liquid fractions were concentrated with rotary evaporator to obtain the crude extracts [25, 26]. Each methanol extract was lyophilized and stored in a moisture-free desiccator until use. The stock solutions were prepared (10.000 μg/mL), and no organic solvents were used to solubilize the extracts, only DMEM medium. Before the assays, the fractions were filtered in Durapore^®^ membrane filter, 0.22 μm.

### Chemical analysis of extracts

The chemical profiles of *C. alba* extracts (CAH21 and CAH24) were analyzed by High-Performance Liquid Chromatography (HPLC) at the Natural Products Laboratory of the Federal University of Tocantins using a SHIMADZU^®^ HPLC system (Kyoto, Japan) consisting of a LC-10ATVp pump, CTO-10A column oven, DGU14A degasser, SCL 10A system controller, Shimadzu SPD-10AT UV-VIS detector and a loop injector with a loop size of 20 μL.

The chromatographic analysis was carried out in gradient conditions using a C-18 reverse phase column (250×4.6 mm, particle size 5 μm, Luna 5 μ C-18). The elution solvent consisted of 0,1% phosphoric acid in water (Solvent A) and 0.1% phosphoric acid in water/acetonitrile/ methanol (54:35:11 v/v) (solvent B). The elution gradient was used as follow: 0-5 min, 0% B; 5-10 min, 30% B, 10−20 min, 40% B, 20-60 min 40% B, 60-70 min 50% B, 70-90 min 60% B, 90-100 min 80% B, 100–110 min 100% B. 110-120 min 100% B. The flow rate was 1.0 mL/min and the detection wavelength was 280 nm. Compounds were identified by comparing sample retention times with 16 available authentic standards (Figure S1) (Sigma – Aldrich, EUA). The concentrations of the identified compounds have not been determined.

### Cells and viruses

Vero Cells (ATCC^®^: CCL81) - from African green monkey kidney (*Cercopithecus aethiops*), available from the Feevale University Molecular Microbiology (LMM) stock (Rio Grande do Sul, Brazil), were cultured in Dulbecco’s modified Eagle medium (DMEM) (Cultilab, Brazil) supplemented with 10% fetal bovine serum (FBS) (Cultilab). Cells were maintained at 37 °C in 5% CO_2_ [27].

Viral isolates used in the experiments were CHIKV # 3 isolate from a Brazilian clinical sample and MAYV BeAr 20290 (GenBank accession no. KT754168). Vero cells were infected with a Multiplicity of Infection (MOI) = 0,01, in a 75 cm^2^ culture flask containing a monolayer of 8.5×10^6^ cells. The infected cells were incubated for 48 h at 37 °C in 5% CO_2_. The supernatant of the cells was collected and kept at -80° C until viral titration [28].

### Viral titration by plaque assay

Titration was performed for each virus, whose experiments were previously standardized in the LMM [29]. The semi-solid medium was prepared with 2X Minimum Essential Medium (MEM) (Cultilab, Brazil), and carboxymethylcellulose (CMC) 1.5% and 1% for titration of CHIKV and MAYV, respectively.

For cell fixation, 4% formaldehyde was added for 30 min, followed by 1X phosphate-buffered saline (PBS, 137.0 mM NaCl, 2.7 mM KCl, 10.0 mM Na_2_HPO_4_, 2.0 mM KH_2_PO_4_, pH 7.4) [30], and 0,2% violet crystal (Synth, BRA) remaining under stirring for 30 min. Then, the violet crystal was removed, and the plates were dried at room temperature. After drying, the formed lysis plaques were counted, and the viral titer was calculated, whose values were expressed in plaque-forming units (PFU/mL).

### Cytotoxicity assay

The cytotoxicity of the plant extracts (CAH21 and CAH24) was assessed through lysosomal viability, using the standardized neutral red incorporation assay at the Feevale University Cytotoxicity Laboratory [31]. For this, 96 well microplates were prepared 24 h in advance, using Vero cells at a concentration of 2×10^5^ cells/well. After the medium was removed, and 150 μL of the serial dilutions of each extract (starting from 1 mg/mL) was added in triplicate. Untreated cells were used as controls, adding an equal volume (150 μL) of DMEM medium supplemented. The plate remained in an incubator at 37°C in 5% CO_2_ for 48 h.

Cell viability was determined considering the absorbance values obtained for each concentration with those obtained for the respective cellular controls (classified as 100% viable). Extract concentration that is toxic to 50% of the cells (CC_50_) was calculated using the dose-response curve, and the experiment was carried out in triplicate with three independent replications.

### Antiviral assays

For the antiviral activity tests, microplates were prepared as previously described. After incubation at 37°C in a humidified environment of 5% CO_2_, the viral suspension (100 PFU/mL) was inoculated, adding only DMEM in the wells corresponding to cell control and cytotoxicity control. After 1 h of incubation, with slow shaking, every 15 min, the respective viral suspensions and the medium (cell control and cytotoxicity control wells) were aspirated, and then the semi-solid medium was added. Different concentrations of plant extracts (40, 60, 80, and 100 µg/mL) were added and designated treatment wells. The highest concentration of the extract was used (100 µg/mL) for the cytotoxicity control. The microplates were incubated at 37 °C under a CO_2_ atmosphere of 5% for 48 h. Thus, they were fixed and stained as previously described. The analysis was carried out in duplicate, and at least three independent experiments were performed, followed by the standardized protocol for each virus.

The percentage of reduction in the number of plaques was calculated as: *A* (%) = 100 – [(*B* x 100) / *C*], where *A* is the % in the number of plaques reduced, *B* is the average number of plaques for treatments and *C* is the viral control number of plaques [32]. Half-maximal inhibitory concentration (IC_50_) with the respective 95% confidence intervals (95% CI), was calculated. From this, the selectivity index (IS) was determined using CC_50_/IC_50_.

### *In silico* analysis of the interaction between molecules of *C. alba* and the nsP2 protease of CHIKV and MAYV

The amino acid sequence of MAYV’s nsP2 protease (GenBank ID: AZM66145.1) was used for the construction of the 3D structure by homology modeling using The Swiss Model Workspace platform (https://swissmodel.expasy.org/) and the Crystal structure of nsP2 protease from the CHIKV [Protein Data Base (PDB) model ID: 4ZTB] as a model. For the CHIKV nsP2 protease structure, the file was obtained directly from the PDB (ID: 3TRK).

*C. alba* molecules – naringin, syringic and chlorogenic acids, vitexin, myricetin, and quercetin, were modeled using Marvin Sketch 18.10 (ChemAxon). Receptors and ligands molecular docking were performed using Autodock Tools 1.5.7 [33], according to the methodology proposed by our colleagues [34]. Docking calculations were performed using AutoDock Vina. Nine docking positions were generated for each ligand, through the interaction with the target proteins returning affinity energy values (Kcal/mol) [35]. Docking position results were analyzed using PyMOL 2.0 [36] and Discovery Studio 4.5 [37] to select the best location for each ligand within the protein target [34].

### Statistical analysis

For all tests, the GraphPad Prism® software (GraphPad Software, San Diego, CA, USA) version 6.0 was used. For cell viability, the results obtained were submitted to the Shapiro-Wilk normality test. For all analyzes, p <0.05 were considered statistically significant.

## Results

### Chemical profiles of *C. alba* CAH21 and CAH24 extracts

High-performance liquid chromatography (HPLC) analysis showed 28 peaks in the CAH21 extract (S2 Fig.), and 21 peaks in the CAH24 extract (S3 Fig.). From the standards used (S1 Fig.), CAH21 extract revealed the presence of a flavonoid – naringin, and CAH24 extract showed the acids: syringic and chlorogenic, and flavonoids: vitexin, myricetin, and quercetin (Table 1).

**Table 1.**
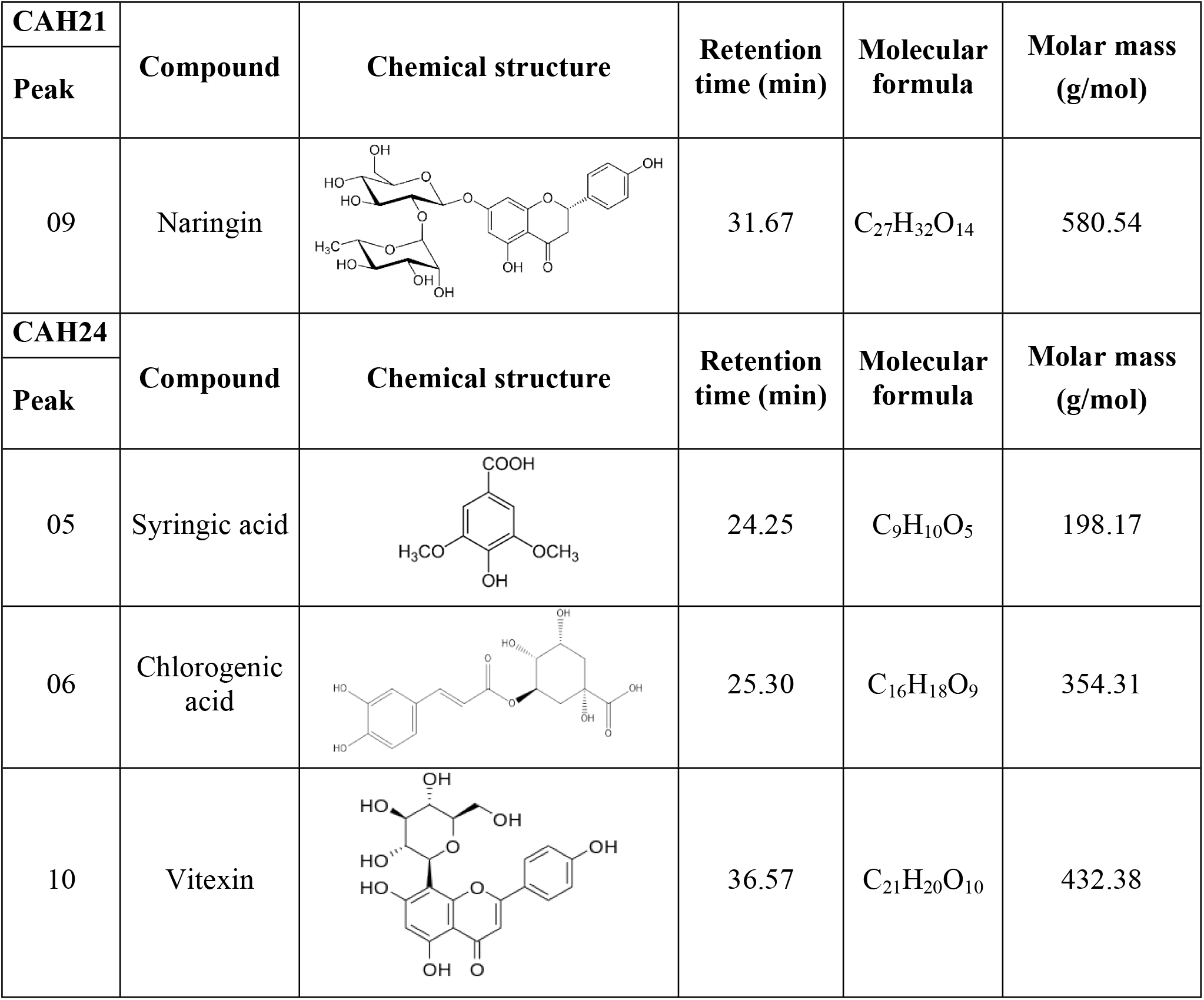
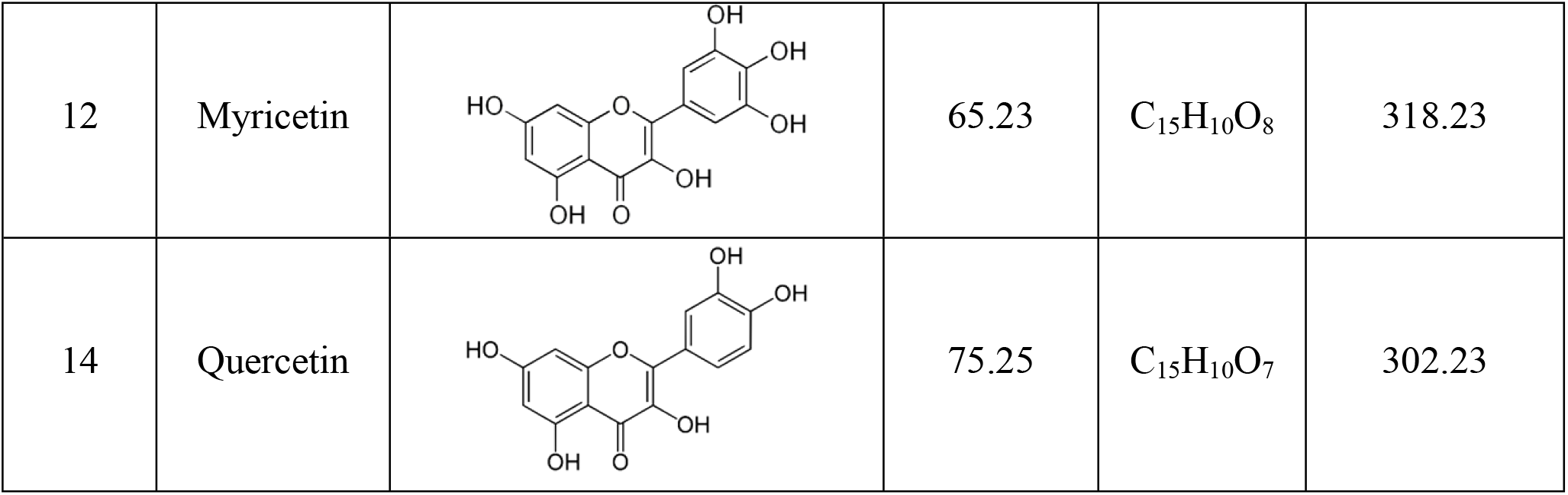
Chemical profile of *C. alba* extracts identified by HPLC.

| CAH21 | Compound | Chemical structure | Retention time (min) | Molecular formula | Molar mass (g/mol) |
| --- | --- | --- | --- | --- | --- |
| Peak |  |  |  |  |  |
| 09 | Naringin |  | 31.67 | C<sub>27</sub>H<sub>32</sub>O<sub>14</sub> | 580.54 |
| CAH24 | Compound | Chemical structure | Retention time (min) | Molecular formula | Molar mass (g/mol) |
| Peak |  |  |  |  |  |
| 05 | Syringic acid |  | 24.25 | C<sub>9</sub>H<sub>10</sub>O<sub>5</sub> | 198.17 |
| 06 | Chlorogenic acid |  | 25.30 | C<sub>16</sub>H<sub>18</sub>O<sub>9</sub> | 354.31 |
| 10 | Vitexin |  | 36.57 | C<sub>21</sub>H<sub>20</sub>O<sub>10</sub> | 432.38 |
| 12 | Myricetin |  | 65.23 | C<sub>15</sub>H<sub>10</sub>O<sub>8</sub> | 318.23 |
| 14 | Quercetin |  | 75.25 | C<sub>15</sub>H<sub>10</sub>O<sub>7</sub> | 302.23 |

### Cytotoxic assay

CAH21 and CAH24 *C. alba* extracts were incubated with Vero cells for 48 h, and the CC_50_ was calculated by nonlinear regression (curve fit). They showed close toxicities to Vero cells in 48 h, and the CC_50_ values were 90 ± 1.4 µg/mL and 100 ± 1.3 µg/mL, respectively (Fig 1 and Table 2). The cytotoxic effects observed were the formation of vacuoles and cell lysis.

**Table 2.** Cytotoxic concentration (CC_50_), selectivity index (SI), and reduction percentage in the plaques number for *C. alba* extracts against CHIKV and MAYV in Vero cells after 48 h.

| Plant extracts | CC<sub>50</sub> (µg/mL)<sup>a</sup> | Concentration (µg/mL) | CHIKV % reduction<sup>b</sup> | Selectivity Index<sup>d</sup> | MAYV % reduction<sup>c</sup> | Selectivity Index<sup>e</sup> |
| --- | --- | --- | --- | --- | --- | --- |
| CAH21 | 89.66± 1.37 | 40 | 47.59±4.62 | 1.66 | 61.62±4.59 | 1.48 |
|  |  | 60 | 73.61±7.98 |  | 79.94±4.39 |  |
|  |  | 80 | 88.03±5.91 |  | 98.14±4.85 |  |
|  |  | 100 | 92.63±3.39 |  | 100±6.40 |  |
| **CAH24** | 100.5± 1.26 | 40 | 60.84±7.49 | 1.76 | 74.35±1.89 | 1.53 |
|  |  | 60 | 79.77±10.13 |  | 83.46±4.69 |  |
|  |  | 80 | 92.33±3.63 |  | 95.79±1.93 |  |
|  |  | 100 | 95.81±0.76 |  | 100±0.00 |  |

<sup>a</sup>CC<sub>50</sub> values represent the means ± SD of three independent experiments (p<0.05).
<sup>b, c</sup>The reported values are mean ± SD of the triplicate assay for each sample.
<sup>d, e</sup>CC<sub>50</sub>/IC<sub>50</sub>.

**Fig 1.**
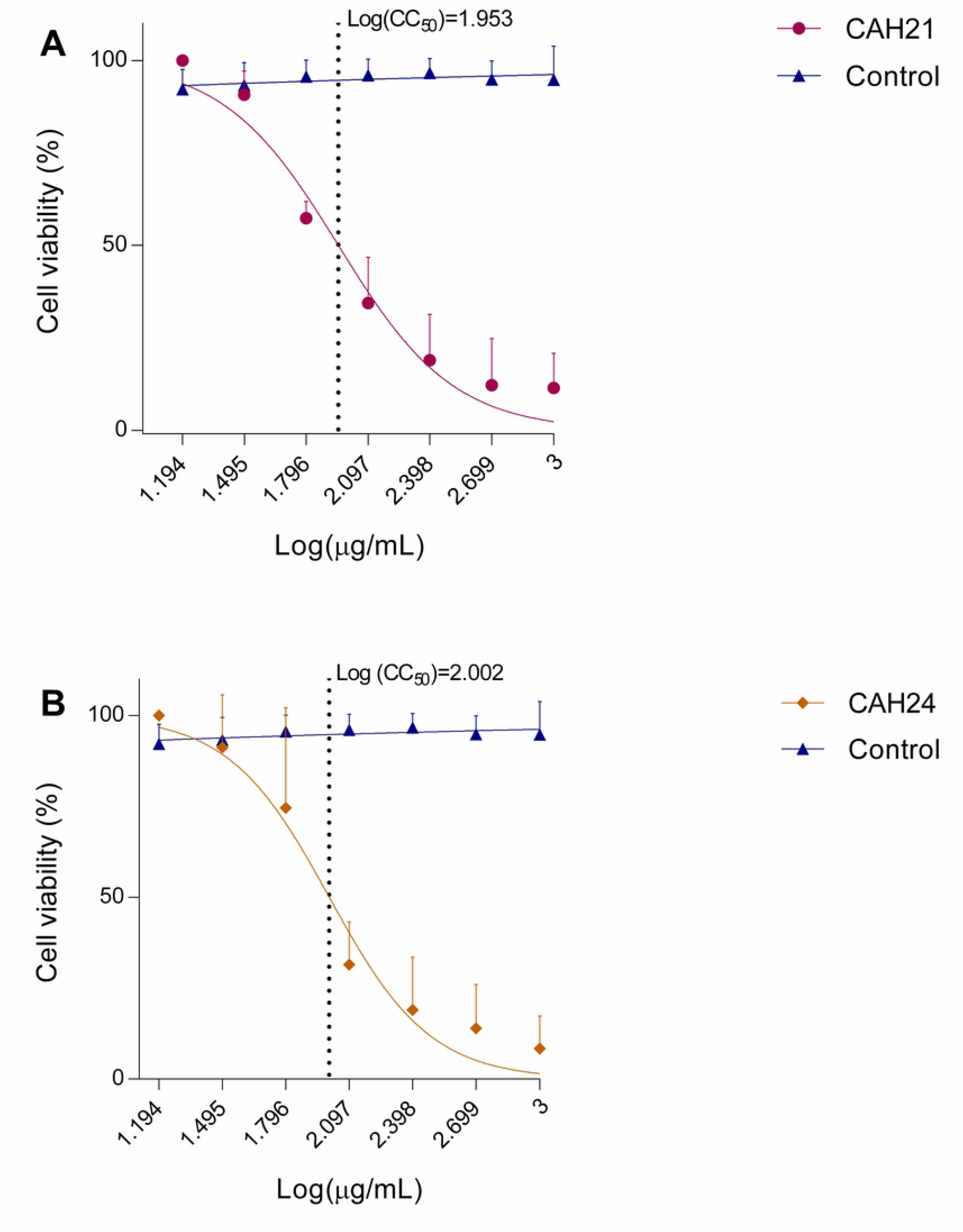
Cytotoxicity assay. Cell viability after treatment using Vero cells, incubated for 48 h, with different concentrations of *C. alba* extracts **(A)** CAH21 and **(B)** CAH24.

### Antiviral activity analysis

Comparing the two extracts, CAH21 showed a less toxic profile for both viruses tested (S4 and S5 Figs.). CHIKV plaque formation reduction, when using the *C. alba* extracts CAH21 and CAH24, was higher than 70% at 60 μg/mL. At 40 μg/mL, CAH21 showed 48% plaque formation reduction (Fig 2 A).

**Fig 2.**
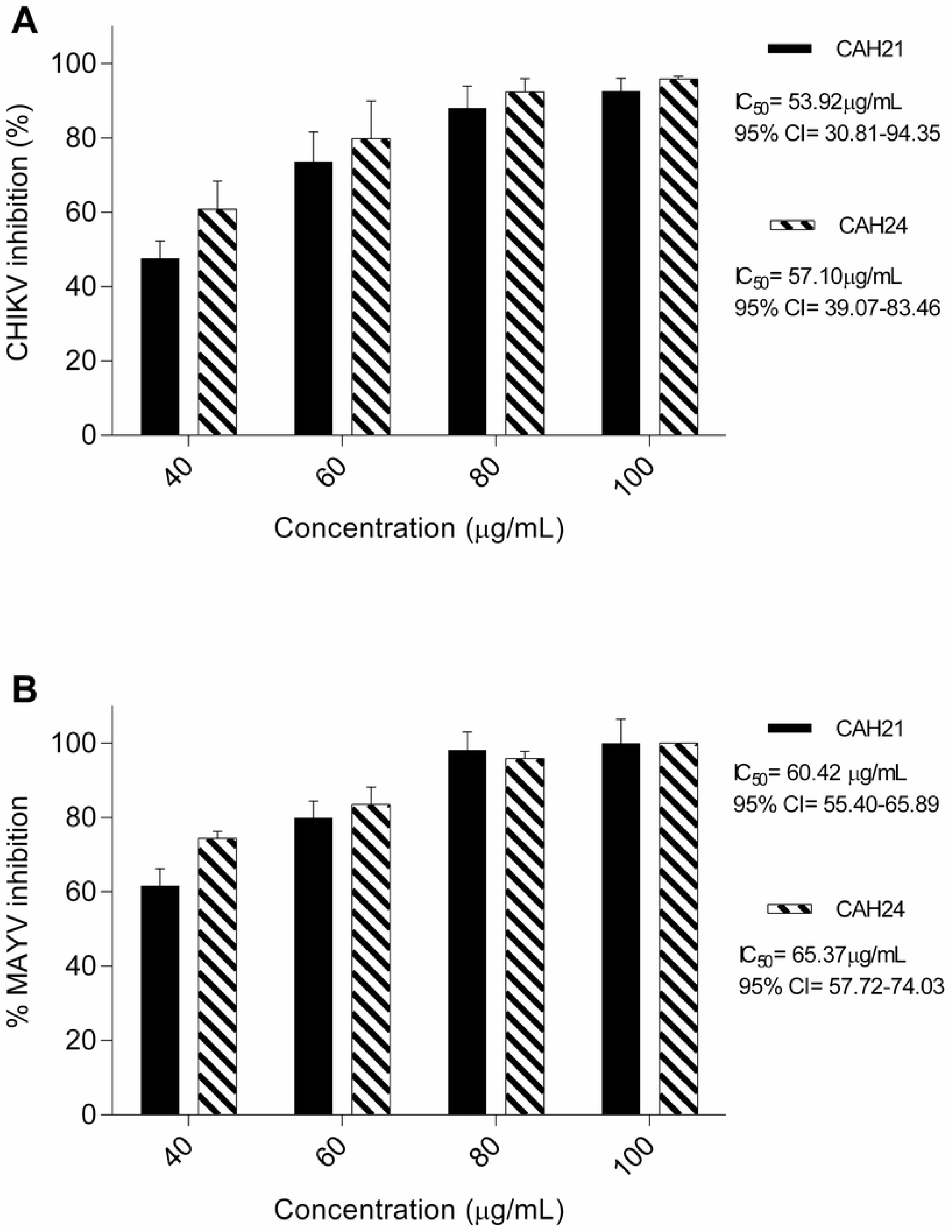
Antiviral assay. Evaluation of the antiviral activity of *C. alba* extracts (CAH21 and CAH24) by reduction percentage in the plaques number of CHIKV (A) and MAYV (B). IC50 = Inhibitory concentration 50% of cells; 95% CI = Confidence intervals of 95%.

The antiviral assays with the CAH24 extract showed a higher reduction in plaque formation than CAH21. For CAH24 extract, 74% and 83% MAYV inhibition was observed at concentrations of 40 μg/mL and 60 μg/mL, respectively (Fig 2 B). Therefore, CAH21 and CAH24 extracts have potential anti-MAYV and anti-CHIKV activity (Table 3).

**Table 3.** Molecular docking results for the complexes formed between the *C. alba* extracts molecules and the nsP2 proteins of CHIKV and MAYV.

| Molecules | Receptor (kcal/mol)<sup>a</sup> |  |
| --- | --- | --- |
|  | nsP2 protease – CHIKV | nsP2 protease – MAYV |
| Chlorogenic acid | -6.7 | -6.8 |
| Myricetin | -7.0 | -7.1 |
| Naringin | -7.9 | -8.5 |
| Quercetin | -7.1 | -7.3 |
| Syringic acid | -4.6 | -4.9 |
| Vitexin | -7.9 | -7.9 |

<sup>a</sup>Affinity energy of Autodock Vina.

### Modeling and validation of the MAYV nsP2 structure

The MAYV’s nsP2 three-dimensional structure was modeled (Fig 4 B) using the CHIKV’s nsP2 as a model (PBD ID: 4ZTB). For the appropriate model construction, amino acid sequences must exhibit very similar structures, as this would imply problems in the functionality of the model. Ideally, this identity should exceed 30% [33], and we found a 67.50% identity about the model used to build the protein three-dimensional structure. About the Ramachandran graph, we obtained 93.08%, and -0.79 to QMEAN analysis (Fig 3 A and B), affirming that the model is located in the most favorable region, close to 0, being an appropriate model [34].

**Fig 3.**
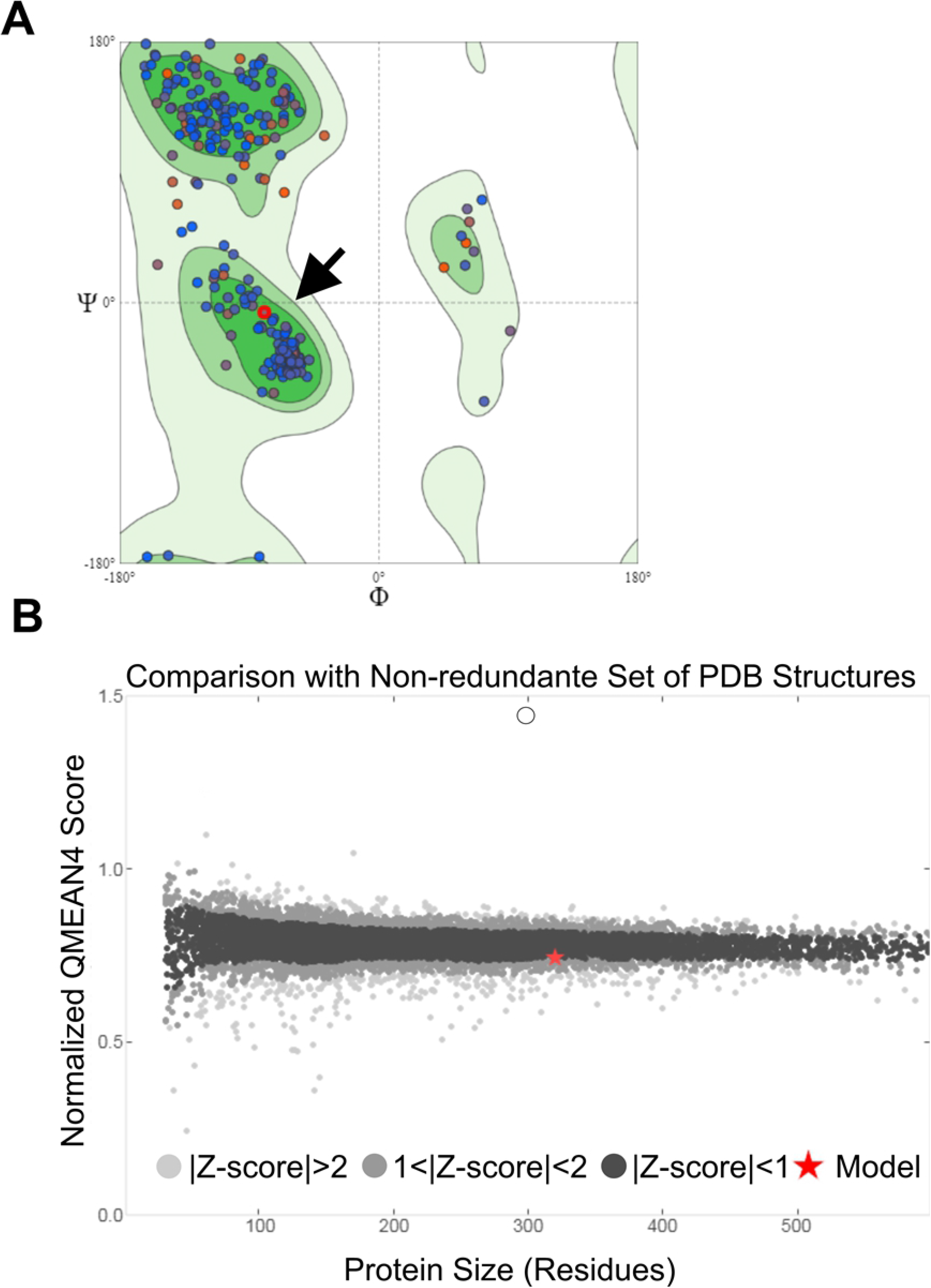
MAYV nsP2 structure modeling and validation. Ramachandran graph for MAYV nsP2 **(A)**. The black arrow indicates the presence of the protein in the most favorable region; QMEAN (Qualitative Model Energy Analysis) value, which describes the quality of the model **(B)**. The red star indicates position in Z-scores presents a highly reliable structure and is within the range of scores typically found for proteins of similar size.

**Fig 4.**
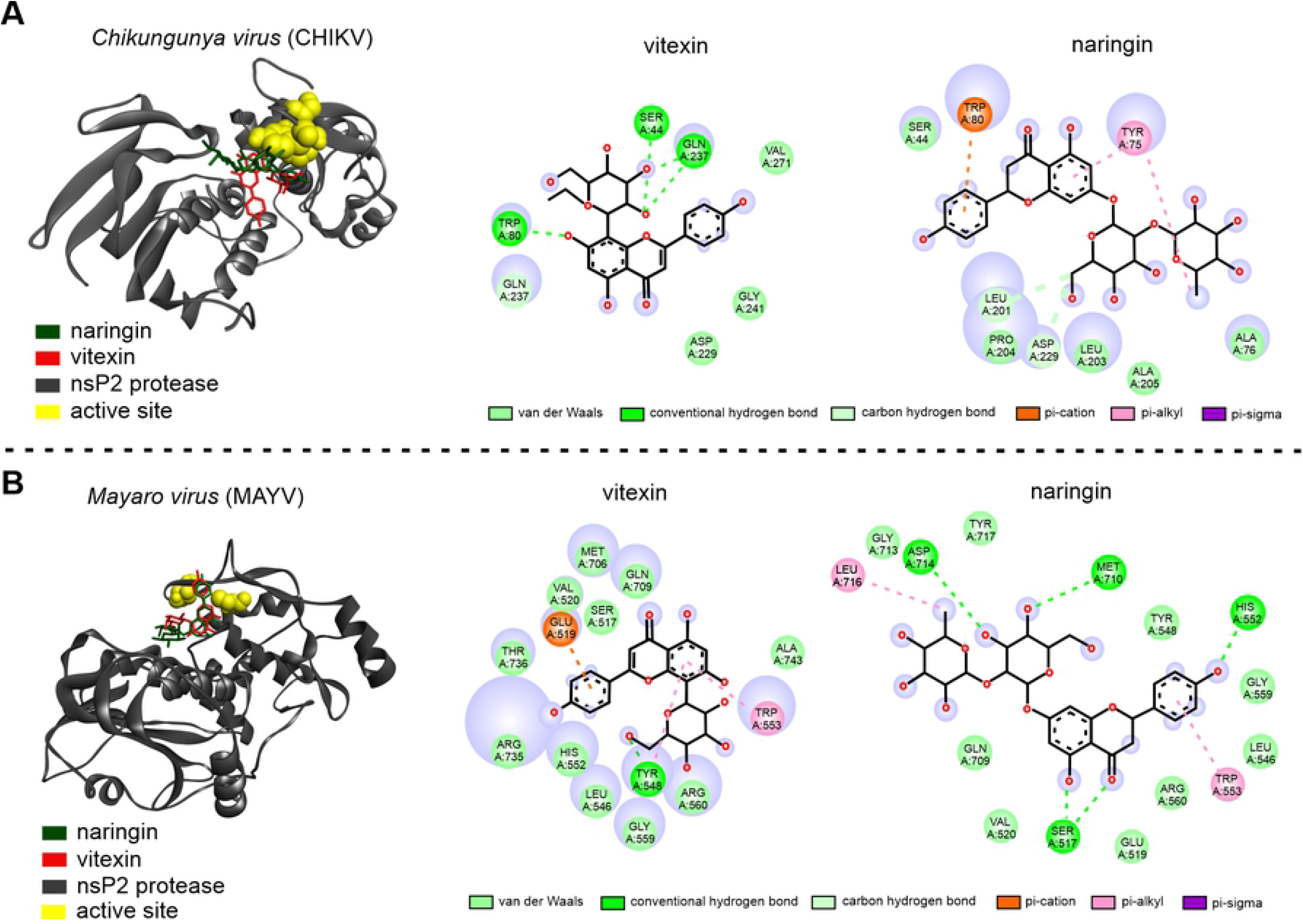
Molecular docking. Complex between nsP2 protease of CHIKV **(A)** and MAYV **(B)** with molecules ligand naringin (green) and vitexin (red), close to the region of the active site (yellow), and 2D interaction map with the amino acids.

### Molecular docking

According to *in silico* analysis, the molecules identified in the *C. alba* extracts can interact with the CHIKV and MAYV nsP2 protein, showing different affinity energies, as described in table 3. Based on the best interaction energies between molecule-receptor, naringin and vitexin showed a strong interaction with nsP2, thus being potential anti-CHIKV and anti-MAYV molecules. The molecular docking using naringin and vitexin with nsP2 of CHIKV and MAYV was used to assess the antiviral potential of these flavonoids, based on the hypothesis that flavonoids may interfere in the viral replication cycle.

Initially, for CHIKV, the nsP2 protease was used as a target (Fig 4 A). Vitexin showed hydrogen bond interactions (SER44, GLN237, and TRP80) with the nsP2 active site. The naringin-CHIKV-nsP2 complex showed van der Waals interactions (SER44, PRO204, LEU203, ALA205, and ALA76), one pi-alkyl interaction and one pi-sigma with TYR75 amino acid, hydrogen carbon interactions (LEU201 and ASP229), and one pi-cation interaction (TRP80) (Fig 4 A). The MAYV nsP2 protease was also used as a target. The vitexin-MAYV-nsP2 showed van der Waals interactions (amino acids MET706, GLN709, SER517, VAL520, THR736, ARG735, HIS552, LEU546, GLY559, ARG560 and ALA743), a carbon hydrogen interaction (TYR548), a pi-alkyl interaction (TRP553), and pi-cation interaction (GLU519) (Fig 4 B). The naringin-MAYV-nsP2 complex showed a total of 15 interactions, with van der Waals bonds (TYR717, GLY713, GLN709, VAL520, GLU519, ARG560, LEU546, GLY559, and TYR548), carbon hydrogen (ASP714, MET710, HIS552, and SER517), and pi-alkyl (TRP553 and LEU716) (Fig 4 B).

## Discussion

In this work, we have showed that *C. alba* methanolic plant extracts have potential anti-CHIKV and anti-MAYV. The cytotoxicity of *C. alba* extracts to Vero cells was determined, establishing a concentration-response curve. An antiviral drug must have activity against the virus without inducing significant toxicity to the host cell, so the first step for antiviral tests is to investigate the cytotoxic activity. Given the crude nature of the tested extracts, the observed cytotoxic effects can be related to the molecules present and theirs concentrations, since the stock solutions was prepared without organic solvents. The higher concentration of metabolites, the greater the cytotoxicity. *In silico* analyzes demonstrate that some of these compounds have a better interaction with the protease nsP2 CHIKV and MAYV, which suggests a model of mechanism.

An extract to be considered with potential antiviral activity will have a reduction of between 50 and 90% in the number of plaques when compared to controls. On the other hand, to be considered antiviral, the extract will need a >90% reduction [38]. The methanolic extract obtained by soxhlet, CAH24, when compared with the maceration method, CAH21, showed higher antiviral potential activity against CHIKV and MAYV. The soxhlet system temperature remains high during the extraction process, higher solubility and diffusivity of the sample is achieved [39]. Therefore, the use of high temperature provided a higher extraction of biologically active antiviral metabolites in *C. alba*, although it also showed a more cytotoxic profile. HPLC analysis of *C. alba* extracts showed 28 (CAH21) and 21 (CAH24) peaks (See S2 and S3 Figs.). However, since we used only 16 known standards molecules (See S1 Fig), it was possible to identify only 6 metabolites: naringin, syringic acid, chlorogenic acid, vitexin, myricetin, and quercetin. The use of different extraction methods and preparation of *C. alba* extracts, aimed to evaluate and compare the profiles obtained in cytotoxicity and antiviral tests, as well as in chromatographic analysis, identifying some different metabolites.

To date, there are no published data on the antiviral activity of *C. alba* extracts against arboviruses. Since we do not know which compound or compounds are responsible for the possible antiviral activity of both extracts, it would be necessary to fractionate the different compounds present in the extract to test their antiviral activity. According to our results, we hypothesize that flavonoids found in *C. alba* could be able to interact with viral replication enzymes, which are described in the literature as having antiviral activity [14, 40, 41], mainly against arboviruses [42-47], and against *influenza viruses* [48], *enteroviruses* [49, 50], and *hepatitis viruses* [51, 52]. Antiviral activity of four flavonoids (quercetin, naringin, hesperetin, and daidzein) was analyzed against *dengue virus*-type 2 (DENV-2) through plaque assay. When it was used after virus adsorption, quercetin showed an IC_50_ of 35.7 µg/mL. Naringin showed an IC_50_ = 168,2 μg/mL before virus adsorption. Daidzein exhibited low activity anti-DENV-2 and hesperetin was not antiviral against DENV-2. Besides, the DENV-2 RNA production level decreased in the presence of 50 µg/mL of quercetin, achieving a 67% viral reduction when compared to the control [46]. The quercetin and the naringenin flavonoids showed antiviral activity against MAYV and DENV-1 to 4, respectively [50, 51].

A screening of methanolic extracts from medicinal plants was performed for anti-DENV activity. The species *Andrographis paniculata* and *Momordica charantia* showed 75 and 50% inhibition in DENV replication, respectively. According to the authors, the flavonoids canferol, quercetin (founds in *M. charantia*), and luteolin were responsible for the antiviral action [53]. Among the aforementioned flavonoids, two were identified in the present study by HPLC: naringin in CAH21 and quercetin in CAH24. Syringic acid identified in CAH24, until the present study, has not been correlated with arboviruses’ antiviral activities, and there are few studies on chlorogenic acid antiviral activity [54]. However, the authors [55] suggested that the antiviral activity (anti-Herpes Simples Virus, HSV) of substances isolated from *Persea americana* species may be due to a synergism between flavonoids and chlorogenic acid.

The flavonoids’ antiviral activity has been related to the inhibition of viral RNA polymerase, adhesion, and entry of viruses into cells [27, 42]. The affinity between the constituents present in the extract and the specific viral proteins depends, in large part, on the types of chemical bonds between these substances and the amino acids of the viral proteins. The separation of nonpolar and polar components from plant extracts may increase the chance of finding highly active antiviral compounds with low cytotoxicity [56]. Therefore, elucidating the antiviral mechanism of *C. alba* extracts requires the identification of their biologically active constituents, allowing the active principle to be used without the interference of other metabolites. The possible antiviral activity found in the *C. alba* extracts can be attributed to the presence of major components, such as flavonoids and/or minor components, unidentified, present in the extract.

In this work, the *in silico* analysis of the possible binding of the metabolites identified in the *C. alba* extracts with the protease nsP2 of CHIKV and MAYV was carried out. Although we have not identified all metabolites of the two methanolic *C. alba* extracts, the molecules that have been detected were used for *in silico* interactions analysis against the CHIKV and MAYV nsP2 proteins. Besides protease activity, nsP2 presents other enzymatic activities during viral replication, such as RNA helicase and RNA triphosphatase – that are responsible for unwinding the duplex RNA and removing the phosphate-5 ‘group from viral RNA during the capping reaction [57]. Molecular interactions analysis by molecular docking, in the active sites of nsP2, for both CHIKV and MAYV, showed that the most common chemical bond found was the van der Waals type. Although the van der Waals type molecular interactions are considered weak, other interactions may happen due to a good correlation between the protein binding site and the ligand structure [58].

CHIKV nsP2 has a high-resolution crystal structure available which facilitates the search for antiviral molecules [59]. To build the MAYV nsP2 crystal structure, the Ramachandran graph is one of the best quality analyses to select experimental structure models. The values report the distribution of the protein torsion angles (φ, Ψ). The reference parameter should be close to 96% [60]. Recently, a flavanone glycoside-naringin that binds to nsP2 protease at nM affinity was identified, via a combination of receptor-based docking and Molecular Dynamics (MD) simulations [61]. The authors submitted the protease-naringin complex to MD simulations, whose objective was to test the stability of the protein-ligand complex. According to the ParDOCK server used, naringin presented a strong connection with the nsP2 active site, which was validated in biomolecular interaction studies, with affinity energy equal to -9.47 kcal/mol, corroborating our results. An *in silico* study of possible molecular interactions between CHIKV nsP3 and three flavonoids potential ligands was conducted. Among of the compounds, the flavonoid baicalin obtained the best affinity binding (−9.8 kcal/mol), showing be like potential inhibitor of CHIKV nsP3 [58].

Until the present study and to our knowledge, this is the first work that shows the interaction between flavonoids and MAYV nsP2 protease *in silico*. However, the antiviral activity of the flavonoid epicatechin isolated from *Salacia crassifolia* against the MAYV capsid protein (protein C) was identified [27]. The activity anti-MAYV and anti-CHIKV of silymarin flavonoid was also confirmed by *in vitro* biologic assays [62, 63]. Plants’ use in Brazilian traditional medicine for the treatment of various diseases is increasing due to the diversity of bioactive constituents present in them, being potential sources of antiviral substances [12, 42]. In this context, the prospection of new drugs of plant origin remains relevant, considering the number of publications demonstrating the antiviral activity of plant derivatives [10], combined with the wide structural diversity found in these substances, in addition to the wide availability in nature [11].

## Conclusion

The *C. alba* roots methanolic extracts showed potential activity anti-CHIKV and anti-MAYV. *In silico* analysis the flavonoids naringin and vitexin showed high ligand affinity to the protease enzyme nsP2 from CHIKV and MAYV, we therefore, hypothesized that these bioligands may inhibit viral replication. As many natural products, *C. alba* roots are a potential source of antiviral compounds for the development of antiviral drugs against viral infections that affect millions of people worldwide.

## Acknowledgments

This work was supported by grants from the National Council of Scientifc and Technological Development (CNPq – process numbers: 313455/2019-8; 427304/2018-0; 308576/2018-7), Tocantins State Foundation for Research Aid (FAPT-SESAU/TO-DECIT/SCTIE/MS_CNPQ/N° 01/2017) and Federal University of Tocantins (PROPESQ) - EDITAL N° 29/2020 PROPESQ and PPGBIOTEC/UFT/GURUPI-Chamada pública para auxílio de tradução e/ou publicação de artigos científicos - EDITAL N° 011/2020.

## Supporting information

**S1 Fig. Authentic standards fingerprint available in High-Performance Liquid Chromatography (HPLC)**. Source: Laboratory of Scientific Instrumentation (LABIC), Federal University of Tocantins.

**S2 Fig. The *fingerprint* obtained by HPLC of the methanolic extract obtained from *C. alba* roots (CAH21)**. Peak 09: Naringin, detected at 280 nm. The separation was performed on a C18 Phenomenex Luna column and gradient system, with mobile phase A 0.1% phosphoric acid in Milli-Q water and mobile phase B 0.1% phosphoric acid in Milli-Q / acetonitrile-water / methanol in the ratio 54:35:11 (v / v).

**S3 Fig. The *fingerprint* obtained by HPLC of the methanolic extract obtained from *C. alba* roots (CAH24)**. Peak 05: syringic acid, 06: chlorogenic acid, 10: vitexin, 12: myricetin, and 14: quercetin, detected at 280 nm. The separation was performed in a C18 Phenomenex Luna column and gradient system, with mobile phase A 0.1% of phosphoric acid in water Milli-Q and mobile phase B, 0.1% phosphoric acid in Milli-Q / acetonitrile-water / methanol in the ratio 54:35:11 (v / v).

**S4 Fig. Plaque assay for CAH21 and CAH24 *C. alba* extracts against CHIKV on Vero cells monolayer for 48 hours**. CC= cellular control; VC= viral control; CC CAH21= cytotoxic control with 100 µg/mL of CAH21, and CC CAH24= cytotoxic control with 100µg/mL of CAH24. Concentrations: 100= 100 µg/mL; 80= 80 µg/mL; 60= 60 µg/mL, and 40= 40 µg/mL.

**S5 Fig. Plaque assay for CAH21 (A) and CAH24 (B) *C. alba* extracts against MAYV on Vero cells monolayer for 48 hours**. CC= cellular control; VC= viral control; CC CAH21= cytotoxic control with 100µg/mL of CAH21, and CC CAH24= cytotoxic control with 100µg/mL of CAH24. Concentrations: 100= 100 µg/mL; 80= 80 µg/mL; 60= 60 µg/mL, and 40= 40 µg/mL.

